# Tunable bicontinuous macroporous cell culture scaffolds via kinetically controlled phase separation

**DOI:** 10.1101/2024.07.18.604180

**Authors:** Oksana Y. Dudaryeva, Lucien Cousin, Leila Krajnovic, Gian Gröbli, Virbin Sapkota, Lauritz Ritter, Dhananjay Deshmukh, Robert W. Style, Riccardo Levato, Céline Labouesse, Mark W. Tibbitt

## Abstract

Three-dimensional (3D) scaffolds enable biological investigations with a more natural cell conformation. However, the porosity of synthetic hydrogels is often limited to the nanometer scale, which confines the movement of 3D encapsulated cells and restricts dynamic cell processes. Precise control of hydrogel porosity across length scales remains a challenge and the development of porous materials that allow cell infiltration, spreading, and migration in a manner more similar to natural ECM environments is desirable. Here, we present a straightforward and reliable method for generating kinetically-controlled macroporous systems using liquid–liquid phase separation between poly(ethylene glycol) (PEG) and dextran. Photopolymerization-induced phase separation resulted in macroporous hydrogels with tunable pore size. Varying light intensity and hydrogel composition controlled polymerization kinetics, time to percolation, and complete gelation, which defined the average pore diameter (Ø = 1– 300 μm) and final gel stiffness of the formed hydrogels. Critically, for biological applications, macroporous hydrogels were prepared from aqueous polymer solutions at physiological pH and temperature using visible light, allowing for direct cell encapsulation. We encapsulated human dermal fibroblasts in a range of macroporous gels with different pore sizes. Porosity improved cell spreading with respect to bulk gels and allowed migration in the porous systems.

## Introduction

Mimicking biological tissues *in vitro* relies on the fabrication of complex and hierarchical three-dimensional (3D) scaffolds with controlled architecture, porosity, dynamics and mechanical gradients.^1–3^ Hydrogels comprise a particularly useful class of materials for 3D culture and tissue engineering.^4–6^ However, many hydrogel platforms offer limited control over both structure and mechanical properties across scales. This is, in part, owing to the nano-scale porosity of many traditional hydrogel networks.^7,8^ Nanoporous hydrogels introduce physical barriers to changes in cell morphology, proliferation, and migration.^9–11^ To accommodate dynamic cell behavior, hydrogels have been engineered with degradable bonds in the network backbone, enabling intrinsic (hydrolytic bonds), externally controlled (photocleavable linkages), or cell-mediated (matrix metalloproteinase-degradable peptides) modification of the network structure.^12–16^ More recently, stress-relaxing, viscoelastic hydrogels have been shown to enable cell spreading and migration via material rearrangement on biologically relevant timescales.^17–19^

A complementary approach to engineer hydrogel-based scaffolds that facilitate cell spreading, migration, and tissue elaboration is to introduce micron-scale pores into the material generating macroporous materials.^20^ Macroporous hydrogels (Ø ∼ 1–100 μm) have been produced using different methods, including the annealing of microparticles, addition of porogens, cryogelation, foam formation, lyophilization, and bioprinting.^21–25^ These methods are either limited in resolution or struggle to reproduce hierarchical and interconnected porosity. That is, when the void space is interconnected throughout the material facilitating transport and exchange through the bulk of the material. Furthermore, some of these approaches require an inclusion and removal of additives, which can limit their use in biomedical applications.^26^ Phase separation in polymeric systems has emerged as a useful tool for the fabrication of macroporous materials without the need for porogens or post-processing; thermal-, solvent-, and polymerization-induced phase separation can generate materials with micrometer-scale porosity.^27–29^

Liquid–liquid phase separation can be induced in a single-phase macromolecular solution by varying external parameters, such as temperature or pH, resulting in a change in the interaction and miscibility of the components.^30,31^ Phase separation can also be triggered by polymerization of one of the components. As described in the Flory-Huggins theory, miscibility in dilute and semi-dilute macromolecular solutions varies with the molecular weight of the components as the critical interaction parameter for phase separation, χc, depends on the degree of polymerization, *N*.^32,33^ Thus, in systems near the coexistence curve, changes in molecular weight of the components during polymerization can trigger phase separation.^34^

In gelling systems, polymerization is accompanied with an increase in the viscosity that diverges upon gelation. This solidification of the polymerizing phase stabilizes and pins the phase-separated topology. This phenomenon has been exploited in the biomaterials community to engineer porous hydrogels for tissue engineering. Macroporous dextran-based hydrogels with interconnected pores were prepared by cross-linking methacrylated dextran in the presence of poly(ethylene glycol) (PEG).^35^ While the formulations conditions provided control over the porosity, the size distribution was broad (Ø ∼ 10–120 μm) in all conditions. This general paradigm was extended using dextran as a macromolecular excluder to fabricate macroporous PEG-based hydrogels for neuronal culture and outgrowth.^28^ Hyaluronic acid (HA) was included as a viscosity modifier to limit the rate of phase coarsening and avoid complete phase separation prior to gelation. Changes in the precursor formulation controlled the final hydrogel porosity (Ø ∼ 0.5–50 μm).

As a complementary approach, phase-separating biomaterials were developed wherein emulsion-based pores were generated through binodal decomposition between GelMA and PEG phases.^36^ More recently, spinodal decomposition of cross-linking poly(vinyl alcohol) and dextran formed interconnected microporous biomaterials, suggesting an inverse correlation between the photopolymerization intensity and the final pore size.^37^ In non-biomedical applications, the kinetics of photopolymerization reactions were exploited to structure phase-separating polymer systems, providing control over the size and interconnectivity of the pores.^38,39^ Irradiation of the photopolymerizable monomer mixture with a UV light intensity gradient resulted in a polymer network with a porosity gradient along the propagation direction of light. These recent investigations suggest that pore formation in phase separating systems can also be controlled carefully by tailoring the rate of polymerization of a gelling phase. Pore growth proceeds as a competition between the rate of phase coarsening and percolation (gelation of the polymerizing phase). In this manner, pore growth can be controlled by the rate of gelation yielding materials with variable porosity from a single formulation. However, extensive investigation of kinetic control in the design of macroporous biomaterials remains unexplored. We hypothesized that the rate of photopolymerization could be used to tune the porosity of macroporous hydrogels during liquid–liquid phase separation.

To demonstrate this, we used photopolymerization-induced liquid–liquid phase separation to fabricate macroporous PEG-based hydrogels. We synthesized norbornene-functionalized star PEG macromers, which were reacted with dithiol linkers through photoinitiated thiol–ene reactions to form the gel phase. Photopolymerized thiol–ene hydrogels have been extensively used in tissue engineering and cell culture, and the polymerization kinetics can be controlled by the irradiation intensity.^40,41^ Furthermore, thiol–ene hydrogels can be functionalized with thiolated bioactive ligands, such as adhesive peptide domains.^42^ To induce phase separation, we polymerized the thiol–ene hydrogels in the presence of dextran (exclusion agent) and hyaluronic acid (HA; viscosity enhancer), yielding materials with micron-scale and interconnected porosity. The polymer fractions of PEG, dextran, and HA provided control over the pore size (range) for fixed polymerization conditions. In addition, we varied the irradiation intensity for a fixed formulation, demonstrating that the average pore sizes can be controlled by the rate of photopolymerization alone, while maintaining a narrow pore size distribution. We explained the scaling of pore size with polymerization kinetics and viscosity by modeling the system based on aggregation of domains of polymer-rich phase that grow and fuse up to a critical size that is pinned at the gel point. Finally, macroporous gels were formed in the presence of primary dermal fibroblasts enabling cell spreading and migration. Overall, this work presents a method to induce and control micron-scale porosity in PEG-based thiol–ene hydrogels as an attractive platform for 3D cell culture and tissue repair.

## Results

### Macroporous PEG-based thiol–ene hydrogels

In single-phase aqueous solutions of PEG and polysaccharides phase separation can be triggered by polymerization of one of the components.^43,44^ Polymerization-induced changes in chain solubility can lead to phase separation via binodal decomposition (metastable state; energy barrier) with local and large fluctuations in concentration or via spinodal decomposition (unstable state with no energy barrier) with global and small fluctuations in concentration (**Figure 1a**).^30,45^ Spinodal decomposition results in the emergence of a bicontinuous network of separating phases that grow in size until complete dephasing. Kinetic trapping of the system is possible if it can be arrested prior to complete dephasing. For example, gelation of one phase prior to complete phase separation of a spinodally decomposing system could lead to an interconnected network of pores that would be beneficial for cell culture applications. Therefore, methods to generate such biomaterials with interconnected porous networks under user-defined control are desirable.

**Figure 1.**
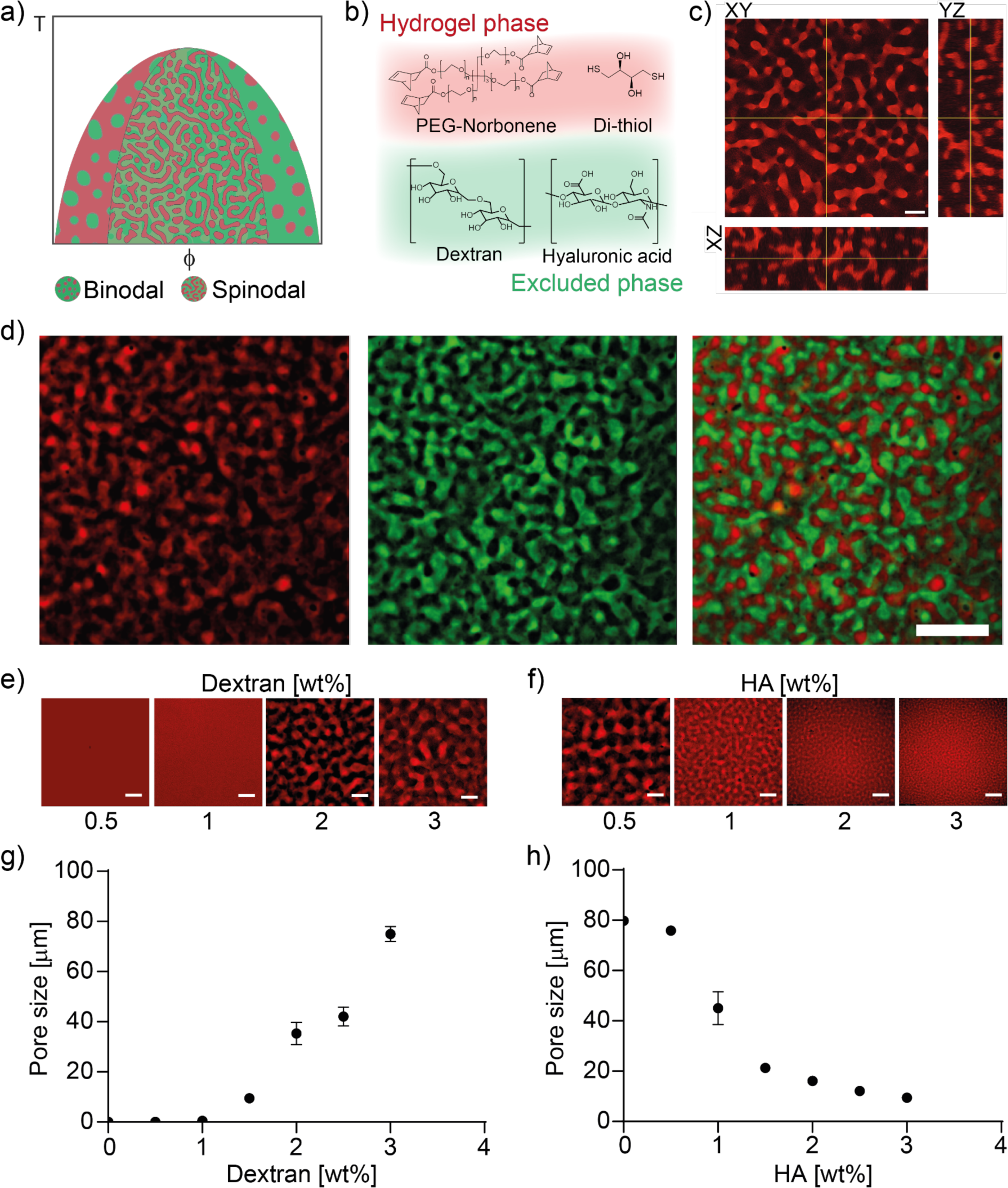
Composition controls phase-separation in polysaccharide containing, photopolymerized thiol–ene hydrogels. a) Simplified schematic of a phase diagram for liquid–liquid phase separation. Phase separation in aqueous solutions of PEG polymers (φ, red) and polysaccharides (green) can be induced through polymerization of the macromolecular components. Phase separation can proceed via binodal or spinodal decomposition mechanisms depending on the mixture composition (φ) and temperature (*T*). b) Macromolecular components of precursor solution for macroporous system include hydrogel phase: 8-arm 20 kDa PEG-norbornene, di-thiol (dithiothreitol; PEG-SH; thiolated peptides); and excluded phase: dextran and hyaluronic acid. c) Irradiation of hydrogel forming solution (λ = 405 nm, *I* = 0.5 mW cm^-2^) triggered thiol–ene polymerization and phase-separation into a bicontinuous network via spinodal decomposition that was arrested by gelation. Hydrogel network phase (red) and polysaccharide phase (black). Scale bar = 50 μm. d) Representative fluorescent images of phase-separated hydrogels (red) perfused with dextran-FITC (green), demonstrating the formation of a bicontinuous and perfusable network. Scale bar =100 μm. e) Representative images of the porosity in 3 wt% PEG hydrogels generated at different dextran concentrations (0.5 – 3 wt%; *I* = 1.0 mW cm^-2^). Scale bar = 50 μm. f) Representative images of the porosity in 3 wt% PEG hydrogels generated at different HA concentrations (0.5 – 3.0 wt%, *I* = 1.0 mW cm^-2^). Scale bar = 50 μm. g) Pore size in 3 wt% PEG hydrogels as a function of dextran concentration (0 – 3 wt%; *I* = 1.0 mW cm^-2^). h) Pore size in 3 wt% PEG hydrogels as a function of HA concentration (0 – 3 wt%; *I* = 1.0 mW cm^-2^).

To generate macroporous hydrogels with controlled porosity, we used a PEG-based thiol–ene system with tunable polymerization kinetics. Hydrogels were fabricated via photopolymerization of norbornene-functionalized 8-arm PEG macromers (*M*n ≈ 20 kDa) and thiol functionalized dithiothreitol (DTT) using equimolar concentrations of functional groups (**Figure 1b**). To introduce porosity, we polymerized thiol–ene hydrogels in the presence of dextran (MW ≈ 100 kDa) that served as an excluding agent and HA (*Mn* ≈ 60 kDa) as a viscosity enhancer to slow down diffusion and enable spinodal decomposition. Upon dissolution, PEG monomers, dextran, HA, and photoinitiator formed a homogeneous mixture. Porous hydrogels formed upon light irradiation (𝜆 = 405 nm, *I* = 1.0 mW cm^-2^) with the photoinitiator lithium phenyl-2,4,6-trimethylbenzoylphosphinate (LAP; **Figure 1c**). Post crosslinking, the pore space was perfused with high molecular weight dextran-FITC. In all cases, the bicontinuous networks were perfusable forming an interconnected network of pores throughout the material volume that enabled transport of dextran-FITC (**Figure 1d**). In order to identify formulations that resulted in the formation of interconnected porosity and to tune the size of the final material porosity, we varied the total polymer fraction and the ratios of dextran and HA to PEG.

The formation of bicontinuous structures was achieved at equal or close to equal weight ratios of PEG to dextran and low concentrations of macromolecules in the solutions ranging from 1.5–4 wt% for both components (**Figure 1e, Figure S1, Movie S1**). The final pore size varied with the total polymer fraction and ratio of PEG to dextran (**Figure 1e,g**). In the absence of dextran, there was no phase separation and PEG hydrogels (3 wt%) formed without the appearance of micron-scale pores (nanoporous). Inclusion of dextran enabled formation of macroporous structures that increased in size from ∼1 to 75.2 ± 2.6 μm over 0 to 3 wt% dextran.

To establish a bicontinuous, phase-separated morphology, we sought to slow the diffusion of the network-forming components, i.e., PEG, without substantially affecting the rate of reaction. This was done as we expected reaction-limited kinetics would favor homogenous network formation prior to pinning via percolation, resulting in a homogenous and nanoporous hydrogel (in the absence of phase separation) or a completely dephased gel (if phase separation did happen). In contrast, we expected that diffusion-limited systems would pin prior to complete phase separation, resulting in distinct, non-overlapping phases. Therefore, HA was included in the precursor formulation to increase the solution viscosity, slowing monomer diffusion, growth of polymer-rich domains, and phase separation to ensure a diffusion-limited system. The inclusion of HA was critical for pore size control, with an inverse correlation between pore size and concentration of HA (**Figure 1f,h**). The pores had diameters of 79.8 ± 1.2 to 9.4 ± 1.1 μm over 0 to 3 wt% HA.

### Tailoring pore structure by varying light intensity

Pore evolution in phase-separating systems depends on the dynamics of phase separation, which controls the growth of polymer-rich domains, and on network percolation, which arrests domain growth and pins them into stationary structures. We hypothesized that by controlling polymerization kinetics we could tune the competition between phase separation and network percolation and, thus, control the size of the pores in the final material. Depending on the polymerization kinetics, the domains can either solidify rapidly producing materials with small-scale porosity or after the system has separated into large-scale porous domains.

The kinetics of thiol–ene polymerization can be controlled by varying the irradiation intensity, which governs the rate of radical generation via cleavage of the photoinitiator. Changing the irradiation intensity (*I* = 1.0–10.0 mW cm^-2^) controlled polymerization kinetics of thiol–ene hydrogels as indicated by the evolution of the storage moduli (G′; **Figure S2**). The evolution of G′ showed a steady increase in the polymerization rate with increasing irradiation intensity. Both the G′/G′′ crossover time, and the time to reach Gmax decreased with increasing irradiation intensity (**Figure 2a, Figure S3**). The former is taken to be the inverse of the rate of polymerization (*Rp*).

**Figure 2.**
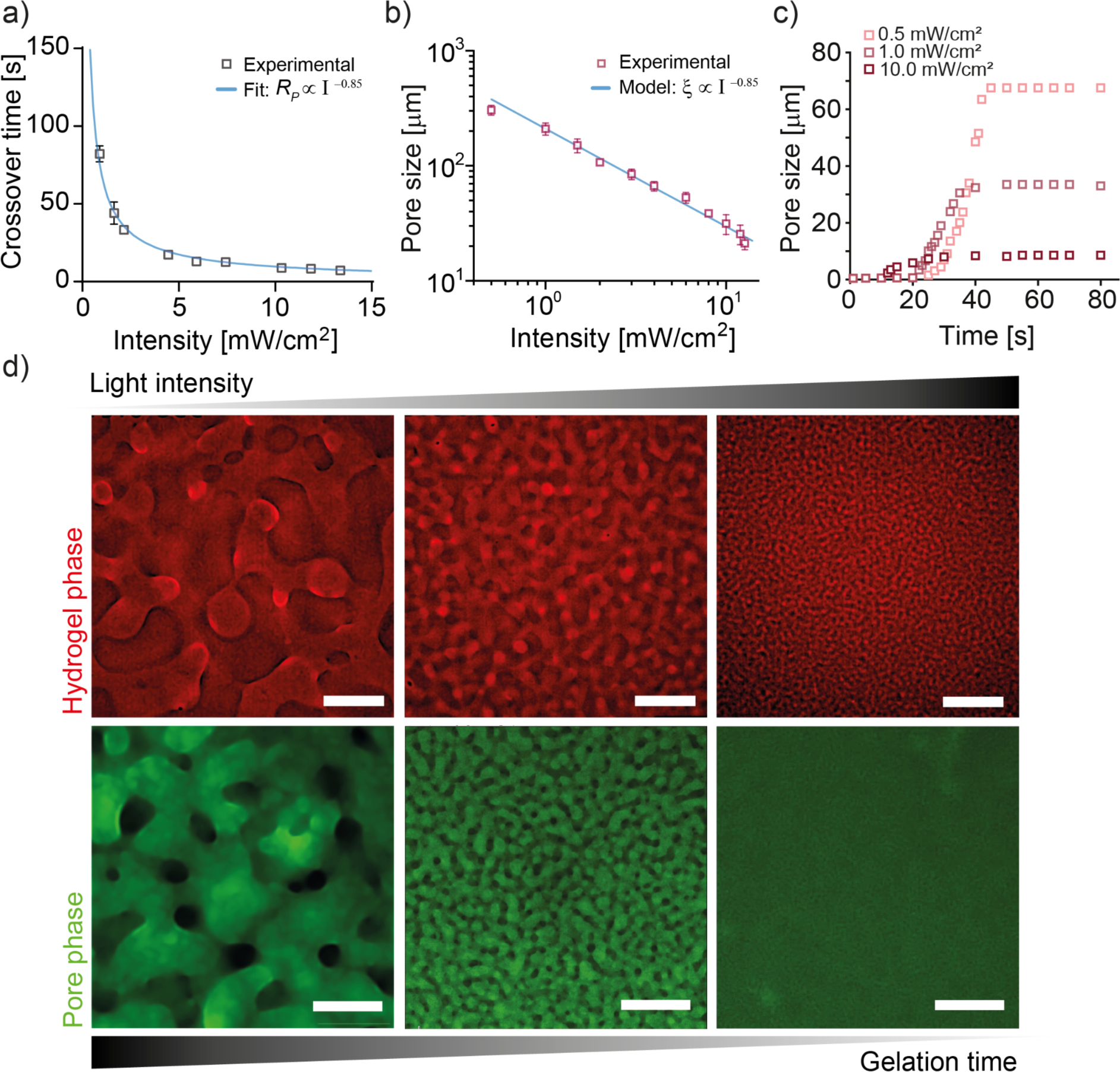
Kinetic control of pore size in macroporous hydrogels. a) In phase-separating, PEG–dextran hydrogels the crossover time, τ*c*, decreased with increasing irradiation intensity, with a scaling based on a power law fit of τ*c* ∼ *I*^-0.85^ or *Rp* ∼ *I*^0.85^. Experimental values are represented as means ± SD (n = 2). b) Pore size in macroporous hydrogels formed in PEG–dextran hydrogels decreased with increasing polymerization intensity, as described by the physical model. Experimental values are represented as means ± SD (n = 3). The solid line represents the scaling law from the physical model (Eq. 3). c) Pore evolution in phase-separating systems at low (0.5 mW cm^2^), medium (1.0 mW cm^-2^), and high (10.0 mW cm^-2^) light intensities (PEG–dextran hydrogels). The light irradiation started at t = 0 s. d) Rhodamine-stained gel phase (red) and dextran-FITC in pore phase (green). Scale bar =100 μm.

Controlling polymerization kinetics using light intensity in PEG–dextran systems controlled the size of the final material porosity. Indeed, upon cross-linking, bicontinuous systems with morphologies formed with characteristic length or pore size, ξ, that decreased with increasing irradiation intensity following a power-law relationship with the percolation time (**Figure 2b,d**). Irradiation at lower light intensities resulted in extended percolation times, which provided longer times for growth and formation of large-scale porosity. High irradiation intensity led to fast percolation of the polymer phase, as indicated by G′/G′′ crossover and rapid immobilization of the formed structures, producing materials with small pore sizes in the range of 10 μm. We confirmed the interconnectivity of the macroporous hydrogels by perfusing them with fluorescein-labeled dextran (MW = 66 kDa), highlighting complete and even perfusion of the pores (**Figure 2d**).

### Polymerization kinetics control macroporous structure

We derived a simple physical model to describe how the average pore size scales with the rate of polymerization (*Rp*, controlled by light intensity) and the solvent viscosity (*ηs*, controlled by HA concentration). The core assumption of the model is that phase separation initiates the formation of polymer-rich domains and polymer-poor domains, where the polymer-rich domains have a characteristic length, ξ. Here, we focus on PEG as the polymer of interest as this comprises the final macroporous network. As we cannot confirm whether the process proceeds with or without an energetic barrier, we consider that the initial shape of the polymer-rich domains is approximately that of a sphere. We allow domains to translate via thermal (Brownian) motion and to encounter each other (**Figure S3**). When two domains come in contact, we assume it is energetically favorable for them to merge. We consider that the merging is driven by the surface tension between the domains and the surrounding medium. Two characteristic times are required to describe the growth of the polymer-rich domains by this behavior: τ*D* the characteristic time for a domain to diffuse and meet another domain, and τm the time for two domains to merge. The parameters τ*D* and τm depend on various parameters, such as the viscosity of the overall medium (*ηs*, influenced by HA concentration), the size of the domains (ξ), and the viscosity of the domains (*ηd,* influenced by the cross-linking density). At the initial stages of spinodal decomposition and photopolymerization, τ*D* is greater than τm allowing spherical domains to coarsen. As the domains grow and become more cross-linked, τm becomes greater than τ*D*. In this regime, additional domains will come in contact with the merging domains during the merging time. This iterative process changes the shape (aspect ratio) of the domains from spheres to more elongated structures (**Figure S3**). As this proceeds, the merging domains will form a bicontinuous network.

We consider that this process determines the characteristic length of the polymer-rich domains, and, thus, the size of the pores of the network, ξ. For this we assumed that the architecture of formed structures remains similar to a large extent until reaching the gelation threshold, where the system is effectively pinned due to the substantial increase in the viscosity of the domains. Based on this physical model, we derived a scaling argument to explain the size of the pores. To develop the scaling parameter, we related τ*D* and τm to relevant system parameters. We assume that the domains reach an equilibrium polymer concentration very fast, based on the assumption that the final gel phase is only twice that of the initial solution given mass conservation. In addition, we neglect the dependence of surface tension on the degree of cross- linking in the domains. We consider that polymer stars do not cross-link outside of domains and that the viscosity change induced by cross-linking inside the domains is much slower than coalescence.

The timescale of merging τm can be obtained as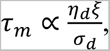, where *η*d, ξ, and *σ*d are viscosity, size, and surface tension of the domains respectively.^46,47^ τ*D* can be approximated by 𝜏_%_ ∝ 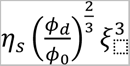 where φd and φ0 are the polymer concentration inside the domains and the initial concentration respectively, and *η*s is the solvent viscosity, based on the distance between domains. Assuming that a bicontinuous network forms when τ*D* ≊ τm, we used this critical point to derive the scaling behavior. In our assumption, *η*d and ξ are the only parameters changing with time during photopolymerization and their evolutions are not correlated a priori. Hence, we derived their evolution as a function of time during photopolymerization.

According to our hypothesis, *η*d can be approximated as *η*d ∝ *η*s · (*R*p*t*)^2(3𝜈-1)^ where *Rp* is the rate of polymerization and ν is the Flory exponent.^48^ By assuming a linear growth in volume, we find that

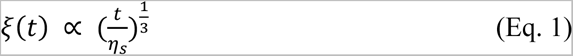

A more detailed derivation of these two scaling relations can be found in the Supporting Information.

Using the expressions for *η*d and ξ as a function of time, we derived the time at which the structures form, i.e., the time at which τ*D* = τm. Using Eq. 1, we calculated the size of the domains at this time:

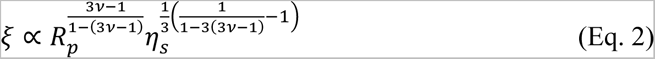

In the case of a theta solvent for the polymer chains, 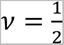 and

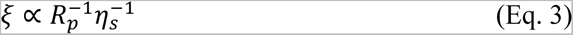

The above framework does not rely on energetics and is consistent with spinodal coarsening theory, which predicts that the length scales in systems undergoing spinodal decomposition coarsen via diffusive, viscous, and, finally, inertial coarsening.^49,50^ In the viscous coarsening regime, which we assume to dominate in the systems with HA, the average length scale should increase linearly with time, i.e., ξ ∝ *t* · *η*s^-1^.^51^ Combining this with the assumption that the time to arrest the system is inversely related to the polymerization rate, *Rp*, implies that the length scale upon arrest should scale as ξ ∝ *Rp*^-1^ · *η*s^-1^.

In each case the predicted scaling follows as ξ ∝ *Rp*^-1^. Given the empirical relationship between *Rp* and *I*, *Rp* ∼ *I*^0.85^ (**Figure 2a**), we arrive at a final scaling relationship between pore size, ξ, and light intensity, *I*, of ξ ∼ *I*^-0.85^, which agrees with the experimental data (**Figure 2b**). The physical model also captures the scaling of pore size, ξ, with solvent viscosity, *ηs*, as ξ ∼ *η*s^-1^, which also agrees with the experimental data wherein we assumed that solvent viscosity was proportional to HA concentration (**Figure S4**). Overall, this model demonstrates how precise control over polymerization kinetics and/or solvent viscosity can be used to tailor feature size, in this case pore diameter, in phase separating macroporous hydrogels.

### Cell spreading and migration

Importantly, bicontinuous structures create an interconnected void space that enables changes in cell morphology and movement in 3D. To demonstrate the effects of scaffold porosity on dynamic cell processes, we encapsulated and cultured human dermal fibroblasts (hDFs) in both nano- and macroporous gels. To enable spreading of the cells that were encapsulated in the gel phase, we introduced proteolytic degradability into our system using thiolated matrix metalloproteinase (MMP)-degradable peptides (KCGPQG↓IWGQCK) for cross-linking our hydrogel system instead of DTT. The thiolated RGD peptides (CRGDS) were covalently coupled to the PEG network to facilitate cell adhesion. The hDFs were encapsulated in nano- and macroporous hydrogels with increasing pore diameters. The cells remained viable under evolution and growth of the spinodal domains and at different porosities proving compatibility of our system with cell encapsulation. The cell spreading in the porous systems was followed for over 7 days and compared with nanoporous hydrogels. The cells exhibited more rapid spreading in macroporous hydrogels than in nanoporous hydrogels in which they remained rounded for over 3 days. In contrast, in macroporous hydrogels cell spreading was observed within a couple of hours and within 3 days cell attained a spindle shaped morphology within the porous space and after 7 days showed an extended spreading (**Figure 3a**). In macroporous hydrogels, cell bodies extended in the pore phase while being anchored to the gel phase (**Figure 3b**). The spreading was faster in the gels with small and medium pore diameters (Ø = 6.0 ± 1.0 and 15.0 ± 1.5 μm pores, respectively) than in gels with large pore diameters (Ø = 45.0 ± 5.6 μm). Furthermore, the cell–cell contacts appeared to be most abundant in the small and medium pore gels. This was shown by the higher number of cell–cell junctions, total cell network area,and the length of cellular structures for small and medium porosity macroporous gels indicating that these systems support a better interaction and interconnectivity of the cells (**Figure 3c-e**). Further, we assessed cell migration within the macroporous materials over 15 h (**Figure 3f-h**). The distances over which encapsulated hDFs migrated in 3D pore space increased with increasing porosity. All porous conditions enabled migration over the distances of > 400 microns within 15 hours (412.3 ± 69.5 μm, 454.3 ± 45.6 μm, and 523.8 ± 69.4 μm in materials with small, medium and large pores respectively; **Figure 3i**). The migration velocity was comparable between all porous conditions (0.5 ± 0.1 μm/min, 0.5 ± 0.03 μm/min, and 0.6 ± 0.1 μm/min in materials with small, medium and large pores respectively; **Figure 3j**).

**Figure 3.**
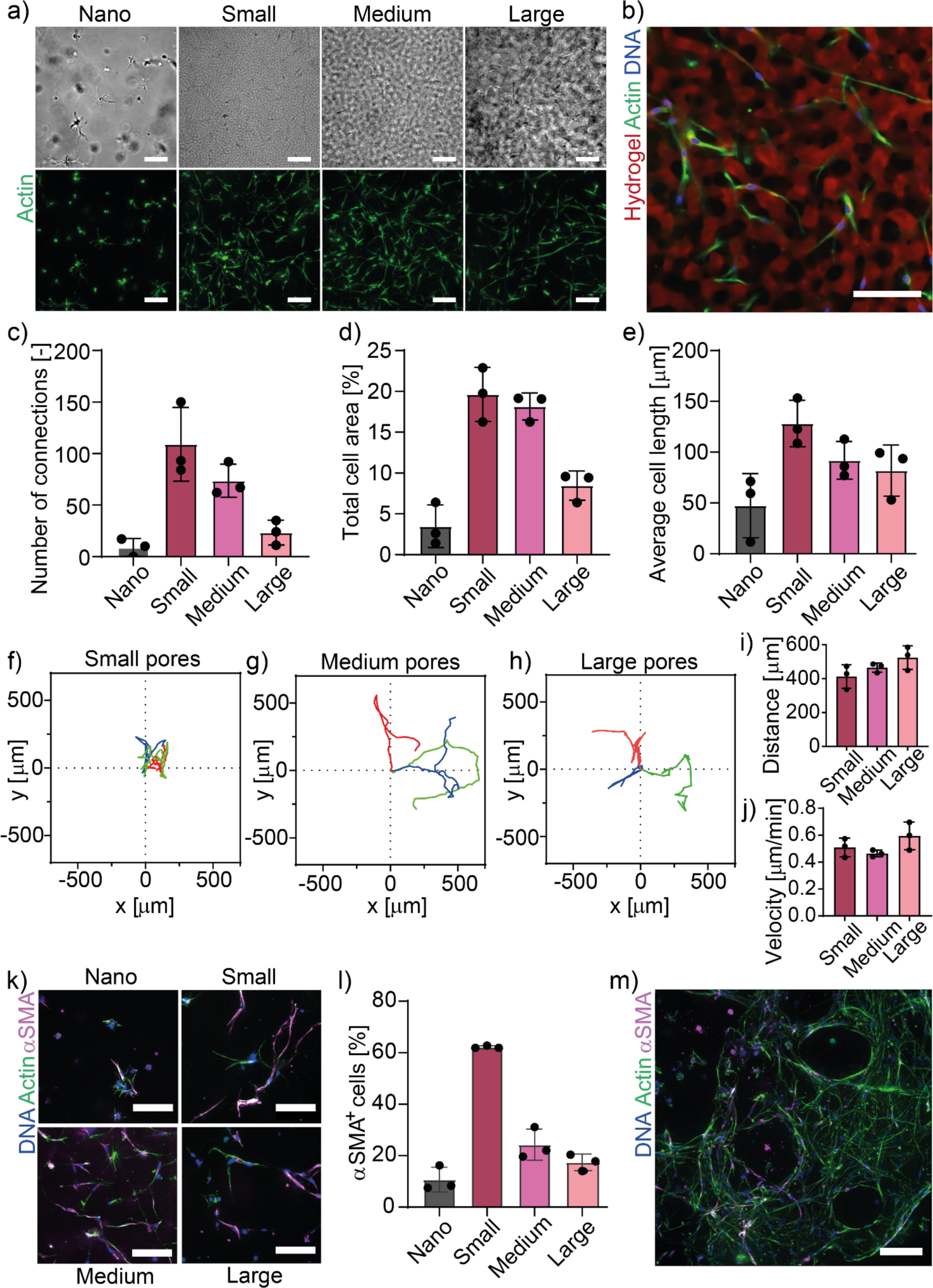
a) Macroporous hydrogels enabled better spreading of the encapsulated hDFs compared to nanoporous gels. Transmitted light images (above) show hydrogel porosity in PEG-MMP hydrogels and fluorescent images (below) show the encapsulated cells 7 days after encapsulation in the corresponding gels with fluorescently stained actin cytoskeleton (green), showing varying degree of cell spreading depending on pore size (nano Ø ∼ 10 nm, small Ø = 6 ± 1.04 μm, medium Ø = 15 ± 1.48 μm, and large Ø = 45 ± 5.56 μm pores). Scale bar = 100 μm. b) Fluorescently labeled cells in microporous PEG-MMP hydrogels with medium porosity (Ø = 15 ± 1.48 μm). Scale bar = 50 μm. c) Percentage of cell area in the macroporous hydrogels with nano, and macroporous hydrogels (AngioTool). d) Number of cell-cell connection in nano, and macroporous gels after 7 days (AngioTool). e) Average spreading hDF length in the nano- and macroporous gels (AngioTool). Trajectories of hDFs migration in macroporous hydrogels with small pores (f), medium pores (g), and large pores (h). i) Distances of hDFs migration in macroporous hydrogels increased with the increasing pore size. j) Velocities of hDFs migration in macroporous hydrogels with small, medium, and large porosities. k) Fluorescent images of cells in nanoporous and macroporous hydrogels at increasing porosity. The morphology of the spread cells and the expression of alpha smooth muscle actin (αSMA) varied between different porosity hydrogels. DAPI (blue, actin (green), αSMA (magenta). Scale bar = 100 μm. l) Expression of αSMA in the nanoporous and macroporous hydrogels at increasing porosity. m) Cells spreading and filling up the macroporous gels with large porosity after 11 days in culture. DAPI (blue, actin (green), αSMA (magenta). Scale bar = 50 μm.

### Pore size influences αSMA expression in cells

In addition to enhancing cell spreading, enabling cell network formation, and facilitating migration, we hypothesized that the porosity of a macroporous 3D scaffold may play a role in cellular mechanotransduction. In 2D culture, the generation of intracellular tension and contractility, is positively correlated with the available spreading area, and there is emerging evidence that the available 3D spreading space, or degree of confinement, has similar effects on cellular mechanotransduction.^52,53^ However, 3D culture systems are often nanoporous hydrogels, which limit cell spreading and the generation of cell contractility in the encapsulated cells.^9^ We hypothesized that cell contractility might be potentiated by enhanced cell spreading and elongation in 3D macroporous substrates. Fibroblast contractility has been found to correlate with *de novo* expression of alpha smooth muscle actin (αSMA) and its incorporation in stress fibers enhancing their stability and myosin light chain activity.^54^

To test whether generation of the contractile fibroblasts increased with the available spreading space in macroporous 3D hydrogels, we cultured hDFs in nano- and macroporous hydrogels with different pore sizes and assessed the number of cells with αSMA-containing stress fibers. After 7 days of culture in nanoporous gels, hDFs appeared more rounded with shorter and less pronounced stress fibers than in gels with macroporosity (**Figure 3k**). The number of αSMA^+^ expressing cells in nanoporous gels was also significantly lower (10.7 ± 4.8%) than in porous hydrogels in which hDFs demonstrated more abundant incorporation of αSMA in stress fibers (above 17% for all conditions) (**Figure 3l**). We hypothesize that the overall increase of αSMA expression in macroporous hydrogels is related to the decreased cell confinement, which enables the generation of more pronounced actin structures and increased cell-cell interconnectivity.

The expression of αSMA varied significantly across the macroporous conditions and exhibited an unexpected pattern. The hydrogels with smallest pores had the highest number of αSMA^+^ cells across all macroporous conditions, while the larger pores, which offered more space, resulted in the lowest numbers of αSMA^+^ cells. The number of αSMA^+^ cells in the hydrogels with the smallest porosity was 62.2 ± 0.5% and decreased at increasing pore size to 24.3 ± 6.1% in medium and to 17.4 ± 3.2% in large pores (**Figure 3l**). The higher αSMA^+^ expression in hydrogels with smaller pores correlated with the cells being more elongated and more interconnected in those hydrogels (**Figure 3k-m**). The cells in the smaller pores tended to organize into more elongated fibrillar structures consisting of multiple cells that may potentiate neighboring cell contractility.

## Discussion

Current 3D cell culture models based on nanoporous synthetic hydrogels limit the generation of functional 3D tissues in vitro in part due to cellular confinement. Notably, confinement hinders essential cellular processes, such as cell growth, elongation, division, and migration. Furthermore, investigating cell mechanobiology, particularly probing cellular responses to stiffness, remains difficult in nanoporous 3D systems because increased stiffness is inherently linked to a more confined microenvironment, making it difficult to isolate the effects of stiffness alone. Many have explored the formation of porous hydrogel materials that would better mimic native ECM porosity and enable biological investigation in 3D; however, specific control over pore size remains challenging.

Phase separation through spinodal decomposition enables fabrication of materials with interconnected micron-scale porosity. Here, we exploited polymerization-induced phase separation in a PEG–dextran solution to fabricate PEG-based hydrogels with engineered porosity for cell culture, building on recent advances in the literature.^28,36,37,55^ We advanced the methods of fabrication of macroporous hydrogels by including kinetic control of the phase separation process using photopolymerization, which provided fine control over the pore size between 1 and 300 μm. Further, we established a model for porosity evolution in thiol–ene PEG–dextran systems, which can be extended to other phase separating systems under kinetic control.

These gels served as a 3D medium for encapsulation and culture of dermal fibroblasts demonstrating that porosity enables improved spreading and communication between the encapsulated cells. In addition, these gels have enabled cell migration over distances > 400 microns, which increased with an increasing pore size. The porosity enabled cell migration with an increased velocity and over longer distances compared to the migration rates reported in the similar or lower stiffness nanoporous systems.^56^ The biological investigation with dermal fibroblasts has shown that the macroporosity allows cell–cell communication and enhanced contractility in the encapsulated cells with respect to nanoporous systems. The increasing pore sizes, however, did not result in a higher contractility. Instead, contractility decreased with increasing pore size. Similar trends have been observed in single cells, which developed more stress fibers and higher actomyosin contractility at intermediate levels of confinement in 3D microniches.^57^ Our results are consistent with these findings, but also suggest that the level of cell-cell interactions is important to cell contractility. This could be further explored by comparing the behavior of different contractile cells (e.g., activated and non-activated fibroblasts), and by varying cell density in our macroporous hydrogels. Instead, contractility decreased with increasing pore size. This intriguing result could be further explored by comparing the behavior of different contractile cells (e.g., activated and non-activated fibroblasts) in macroporous materials with different pore sizes. Our observations suggest that the level of cell-cell interactions could be important, which could be tested by varying cell seeding density. Other potential factors in the contractility scaling could be the relative size of pores with respect to the cell width, and the shape rather than the size of the pores.

## Conclusion

This study demonstrates how control over polymerization kinetics and precursor formulation can be leveraged to precisely tune the final porosity in phase-separated, macroporous hydrogels. This approach provides a method to engineer porous materials for biomedical research with defined characteristics. We demonstrate that macroporous hydrogels fabricated via phase separation offer a superior 3D cell culture environment compared with nanoporous hydrogels. The engineered macroporosity overcomes limitations of nanoporous hydrogels facilitating improved cell spreading, communication, migration, and even regulates contractility, mimicking features of the natural extracellular matrix. This novel approach provides a valuable tool for researchers to study cell behavior and function in a more physiologically relevant 3D setting, overcoming limitations of current in vitro models.

## Author Contributions

The project was conceived and designed by O.Y.D., L.C., and M.W.T. O.Y.D., L.C., R.L., C.L., and M.W.T. designed experiments. O.Y.D., L.C., L.K., G.G., V.S., L.R., R.S., D.D. and C.L., carried out the experiments and analyzed data. O.Y.D, L.C., C.L., and M.W.T. wrote the manuscript. All authors have approved the final version of the manuscript.

## Acknowledgements

The authors would like to thank ScopeM (ETH Zurich) for assisting designing imaging and image analysis strategies; and the SKINTEGRITY.CH cell bank for providing hDFs for the cell investigations. This work was supported by the Swiss National Science Foundation (SNSF) Sinergia Project No. CRSII5_213498, the Helmut Horten Stiftung, and the Open ETH project SKINTEGRITY.CH.

## Methods

### Synthesis of norbornene-functionalized PEG

8-arm PEG amine (Mn ∼ 20 kDa; 4 g, 0.2 mmol PEG, 1.6 mmol NH2, 1 eq. NH2; JenKem USA) was dissolved in anhydrous dimethylformamide (DMF; 5 mL; Sigma-Aldrich) and purged with argon. N,N-Diisopropylethylamine (DIPEA; 1.1 mL, 6.4 mmol, 4 eq.; Sigma Aldrich) was added to the PEG solution followed by the addition of 1- [bis(dimethylamino)methylene]-1H-1,2,3-triazolo[4,5-b]pyridinium 3-oxide hexafluorophosphate] (HATU; 2.43 g, 6.4 mmol, 2 eq.; Sigma-Aldrich). Next, 5-norbornene- 2-carboxylic acid (0.78 ml, 6.4 mmol, 4 eq.; Sigma-Aldrich) was added to the solution and the reaction was stirred overnight at room temperature (RT) under argon. The product was precipitated twice in diethyl ether (4 °C) and the precipitated polymer product was recovered and dialyzed in a dialysis membrane (MWCO 1000 g mol^−1^; Spectrum Laboratories) against dH2O for three days. The aqueous polymer solution was lyophilized obtaining the product in the form of a white amorphous solid. Functionalization of the 8-arm PEG with norbornene was determined to be above 95% via ^1^H NMR in CD2Cl2 by comparing the integrated areas under the peaks of the norbornene vinyl protons (δ = 6.0–6.3, m, 2H) and the PEG ether protons (δ = 3.5–3.9, m, 96H).

### LAP synthesis

Lithium phenyl-2,4,6-trimethylbenzoylphosphinate (LAP) was synthesized according to preveoiusly published synthesis procedure.^58^ In short, 2,4,6-trimethylbenzoylchloride (3.2 g, 18 mmol) was added dropwise to an equimolar amount of continuously stirred dimethyl phenylphosphonite (3.1 g; 18 mmol) at RT and under argon. The reaction was stirred overnight under argon. Lithium bromide (6.1 g; 72 mmol) was dissolved in 100 mL of 2-butanone and added to the reaction mixture. The reaction was heated to 50 °C for 10 min until formation of solid precipitate. The reaction mixture was cooled to RT for 3 h. The precipitate was recovered through filtration and washed 3 times with 2-butanone (50 mL) to remove unreacted lithium bromide and dried under vacuum. The product was recovered in near quantitative yield. ^1^H NMR in D2O: 7.57 (m, 2H), 7.42 (m, 1H), 7.33 (m, 2H), 6.74 (s, 2H), 2.09 (s, 3H), 1.88 (s, 6H).

### Sigmacote coating

To coat a substrate with Sigmacote, the glass slide substrate was cleaned in strong soap (SDS, Sigma-Aldrich) and washed thoroughly with water, ethanol, and acetone. After air drying, the glass slide was placed in a staining container and submerged in Sigmacote (Sigma-Aldrich) for 5 min. The container was sealed with a lid to avoid evaporation of Sigmacote. Subsequently,

Sigmacote was removed and the surface and the surface was washed twice with dH2O and air dried. The substrate was oven dried at 100 °C for 30 min to produce a durable coating.

### Bulk hydrogel synthesis

20k 8-arm PEG hydrogels (3 wt%) were prepared using 8-arm PEG-NB (30 mg; 12 mM NB), freshly prepared DTT (8.5 mg; 12 mM SH), LAP (2.5 mg; 0.25 mM), CRGDS (1 mM), and dH2O.

Two Sigmacote-coated glass slides were separated by a silicon spacer (0.5 mm). The hydrogel precursor solution was injected between the two glass slides and subsequently polymerized under exposure to UV light (λ = 405 nm; *I* = 0.5–15.0 mW cm^-2^) for 1 to 5 min depending on the gelation time of the hydrogel determined during rheological testing. After careful removal of the gel from the Sigmacote-coated glass slides, the gels were immersed in PBS or culture medium and stored in the incubator to equilibrate at 37 °C. The samples were then stored in the incubator for further cell culture without shaking.

### Porous hydrogel synthesis

Porous PEG hydrogels (3 wt%) were prepared with *E* ∼ 1 kPa. For 1 mL hydrogel forming solution, 8-arm 20 kDa PEG-NB (30 mg; 12 mM NB) was mixed with freshly prepared DTT (8.5 mg; 6 mM; 12 mM SH), LAP (2.5 mg, 0.25 mM), CRGDS (1 mM), and hyaluronic acid (HA) as a viscosity enhancer (10 mg, MW = 60 kDa), and dextran (30 mg, MW ≈ 100 kDa; Sigma-Aldrich). The mixture was diluted in HEPES buffer (0.2 M, pH 7.4) to a volume of 1 mL.

Two Sigmacote coated glass slides were separated by a silicon spacer (0.5 mm). The hydrogel precursor solution (20 μL) was injected between the two glass slides and subsequently polymerized under exposure to UV light to attain different porosities (ThorLabs LED, λ = 405 nm; *I* = 0.5–15.0 mW cm^-2^) for 1 to 5 min depending on the gelation time of the hydrogel determined during rheological testing. After careful removal of the gel from the Sigmacote treated glass slides, the gels were immersed in PBS or culture medium and stored in the incubator to equilibrate at 37 °C.

### Rheological characterization

The cross-linking kinetics and mechanical properties of the bulk and porous PEG hydrogels were quantified using a shear rheometer (MCR 502; Anton Paar). The hydrogel properties were assessed in situ or using pre-made hydrogels after equilibration. For in situ rheometry, the hydrogel precursor solution was loaded between an 8 mm parallel plate geometry and a transparent bottom plate with a gap of 0.5 mm. The gel forming solution was cross-linked upon exposure to collimated UV light (λ = 365 nm, *I* = 0.5–15.0 mW cm^-2^; M365L3-C1, ThorLabs). For pre-made hydrogels, samples were loaded between a Peltier plate and an 8 mm geometry with sandblasted surface. To minimize evaporation of water from the gel surface during measurement, a tissue soaked with HEPES buffer (0.2 M, pH 7.4) was placed on the glass along metal ring of the glass plate to surround the sample and a solvent trap was placed over the measurement geometry. Storage (*G′*) and loss (*G″*) moduli were measured over time with oscillatory strain measurements at γ = 1% amplitude and frequency of 1 Hz. All measurements were carried out in the linear viscoelastic regime for the formed gels. The Young’s modulus (*E*) was estimated using the relation between shear and Young’s moduli for isotropic and homogeneous materials, *E* = 2*G′*(1+ν) where ν is the Poisson’s ratio for the material. For the PEG-based materials tested, ν was assumed to be 0.5.^59^ All measurements were performed at 25 °C.

### Visualization of hydrogel porosity

In order to visualize the hydrogel structure, the hydrogel was labeled with acryloxyethyl thiocarbamoyl Rhodamine B (Sigma-Aldrich). The acrylated Rhodamine B was dissolved in dH2O at a concentration of 1 mg mL^-1^. From this stock solution, 50 µL was added to 1 ml of hydrogel precursor solution (0.005 wt%). The hydrogel was polymerized under each Teflon mold and immersed in PBS for 30 min to swell out all unreacted Rhodamine dye. Subsequently, the pattern was imaged with a Leica Thunder microscope (objective 20x, NA = 0.4).

### Image processing and measurement of pore size

For each gel, a z-stack was taken with a Leica Thunder microscope. The average pore size for each gel was determined by measuring manually the size of 15 pores per image. From these measurements the average and standard deviation were computed. In the case where less than 15 pores were visible on the image, the size of all pores was calculated and the average and standard deviation were computed in the same way.

### Derivation of the scaling laws

In the following derivations, only scaling arguments were used. Scaling laws were denoted using the symbol ∝ instead of =. The terms on the left and right-hand side of the sign ∝ are proportional: *A* ∝ *B* means that there is another quantity *K*, such that *A* = *K* × *B*. In deriving the following expressions, the focus was on understanding the influence of the quantities that were varied experimentally, for example, the viscosity of the solvent *η*s, the rate of polymerization *R*p, and the time after the start of the reaction *t*. All other physical quantities were included in the proportionality constant relating the terms on the left- and right-hand side of the equations. Hence, this proportionality constant contains physical units. Because this constant was not explicitly mentioned in the scaling laws, the units of the terms on the left- and right-hand side of the equations may not appear consistent.

### Viscosity of the PEG-rich domains

The intrinsic viscosity of a solution of unentangled polymers can be written, according to the Zimm model, as *η*s ∝ b^3^*N*av/*M*0 · N^(3𝜈-1)^.^48^ In this expression, *b* is the length of a monomer, *N*av is Avogadro’s number, *M*0 the molecular weight of a monomer, 𝑁 the degree of polymerization, and ν the Flory exponent. Considering that the viscosity of the solvent is negligible compared to the viscosity of the polymer solution, one can write *η*d ∝ *η*s · N^(3ν-1)^.

The average degree of polymerization can be approximated using the Carothers equation.^60^ In the limit of small extent of reaction, one can use a first-order Taylor series expansion and write *N* ∝ p^2^, where p is the extent of reaction, p = *R*p · *t*. The presence of p^2^ instead of p in the Carothers equation originates from the fact that each end of the small linker DTT has to be reacted in order to form an effective link, which occurs with probability p^2^. Combining the expression of *N* as a function of *R*p and *t*, and the expression of the intrinsic viscosity, we arrive at *η*d ∝ *η*s · (*R*p*t*)^2(3𝜈-1)^.

### Growth of the PEG-rich domains

Assuming a linear growth in volume, we can write ξ(*t*)^3^ = V(*t*) = C × *t*, where C is a factor that depends on the parameters of the system but not on time. Notably, one can expect C to be linearly related to the coefficient of diffusion of the 8-arm polymer stars in the solution: C ∝ *D*. Using Einstein’s relation D = *k*BT/6π*η*sr, one finds ξ(*t*) ∝ (t/*η*s)^1/3^.

### Cell encapsulation and culture Cell culture

The cell encapsulation experiments were conducted with a human foreskin fibroblasts cell line (Lonza, NHDF NEO 18TL). The cells were cultured in Dulbecco’s modified eagle medium with 4.5 g L^-1^ D-glucose (DMEM (1x) + GlutaMAX, Sigma-Aldrich, St. Louis, USA), supplemented with 10% FBS (Fetal Bovine Serum, GibCo, Waltham, USA) and 1% Penicillin-Streptomycin (Pen/Strep, GibCo, Waltham, USA). The cells were incubated under standard cell culture conditions at 37 °C in a 5% CO2 atmosphere. The medium was changed every two days until the cells were nearly confluent and were passaged, frozen or used for cell encapsulation experiments. For passaging, the cells were washed with PBS, followed by treatment with 0.25% Trypsin-EDTA (Gibco, Waltham, USA) for 2 to 5 min at 37 °C to facilitate detachment from the cell culture flask. The detached cell suspension was diluted with medium centrifuged at 300 rcf for 5 min. The supernatant was aspirated and cells were resuspended in 1 mL of fresh medium. 10 µL of this suspension was mixed with 10 µL of 0.4% trypan blue (NanoEntek, Seoul, Korea) and the mixture pipetted into an EVE cell counting slide (NanoEntek, Seoul, Korea) for electronic cell counting with an EVE Automatic Cell Counter (NanoEntek, Seoul, Korea). The average of the two inlets of the cell counting slide was calculated to get the approximate cell number per mL of cell suspension. The result was used to further calculate the desired volume of the cell suspension needed for encapsulation experiments.

### Cell encapsulation

For 3D cell culture encapsulation, PEG hydrogels (3 wt%) were prepared with MMP cleavable crosslinker (KCGPQG↓IWGQCK; MW = 1304.55 g mol^-^¹; 6 mM; 12 mM SH) instead of DTT, additionally CRGDS peptide (1 mM, GenScript) was added for cell adhesion. The cells were diluted in HEPES buffer (0.2 M, pH 7.4) prior to mixing with the rest of hydrogel components. The hydrogel precursor solution with cells was stored on ice.

For sterilization, glass slides were immersed in ethanol before transferring to the biosafety cabinet for drying. Two Sigmacote-coated glass slides (coated according to manufacturer’s protocol) were separated by a silicon spacer (0.5 mm). 10 μL of hydrogel precursor solution was injected between the two glass slides and subsequently polymerized under exposure to UV light (λ = 405 nm; *I* = 4, 1.2, 0.5 mW cm^-2^ for generation of small, medium and large porosity respectively). The irradiation time to achieve full gelation was determined rheologically (t0.5 = 9 min, t1.2 = 6 min, t4 = 5 min). After careful removal of the gel from the Sigmacote treated glass slides, the gels were immersed in culture medium and stored in the incubator to equilibrate at 37 °C. The samples were then stored in the incubator for the further cell culture without shaking. The medium was exchanged three times every 20 min after encapsulation to accelerate swelling out HA and dextran. A final medium exchange was done at the end of the day. Afterwards the medium was exchanged every 2–3 days.

### Cell migration

Cells were encapsulated as previously described. The cell-laden hydrogels were superficially dried with a sterile Kimwipe tissue (Kimberly-Clark Professional) and cross-linked onto the surface of a 24-well plate using bulk PEG hydrogel precursor solution consisting of 3 wt%, 6 mM MMP, and 0.2 wt% LAP was pipetted onto a well of a 24-well plate (7.5 µL) that was placed between cell-laden hydrogel and the surface of the well. The precursor mixture glued the gel to the surface by crosslinking it using UV irradiation for 2 min with a light intensity of 10 mW cm^-2^ (λ = 405 nm, ThorLabs, M405LP1-C1). The imaging was performed on THUNDER Leica Live Cell microscope with environmental control (37 °C with 5% CO2). The time lapses were recorded over 9 h and the images were collected every 15 min. The migration tracks in x-y plane were recorded using manual tracking ImageJ plug-in.

