## Supporting Information for "Tunable bicontinuous macroporous cell culture scaffolds via kinetically controlled phase separation"

**Movie S1**. Formation of macroporous domains upon exposure to I = 4.0 mW $cm^{-2}$ with 3 wt% PEG-NB, 3 wt% dextran, 1 wt% HA, and 0.25 mM DTT. The PEG phase is tagged with rhodamine. The timelapse shows the onset of phase separation and the formation of the gel phase (red) and the pore phase (black). Time step = 0.5 s.

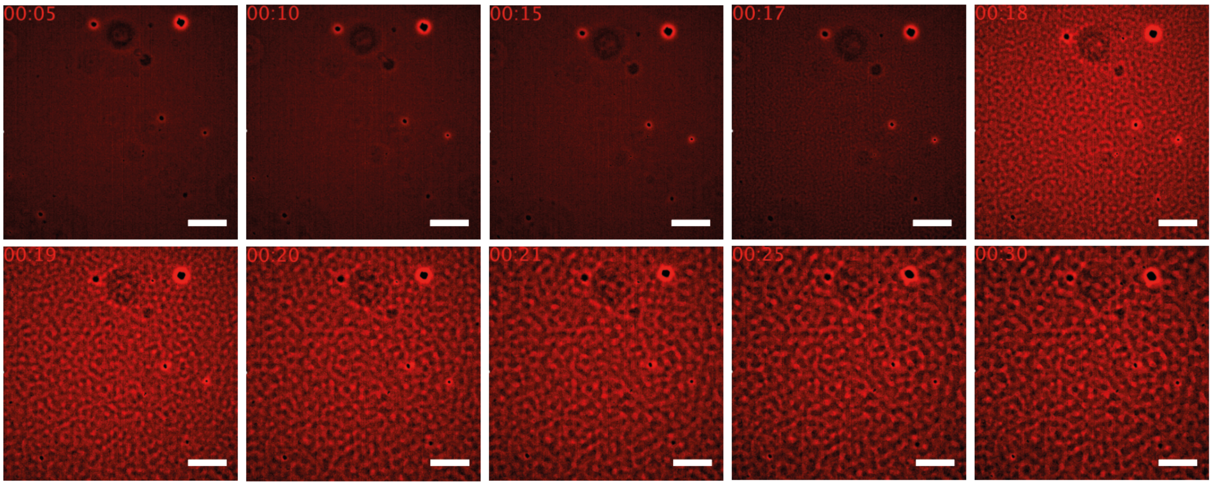


**Figure S1**. Porosity evolution of PEG-dextran system (PEG-NB (3wt%), dextran (3wt%), HA (1wt%) DTT 0.25mM) upon exposure to I = 4.0 mW $cm^{-2}$. Snapshots corresponding to **Movie S1.**

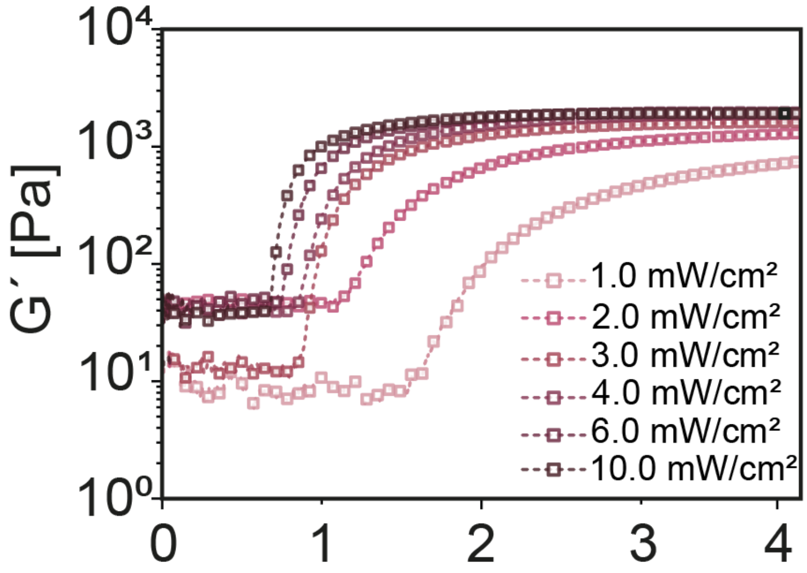


**Figure S2.** Polymerization kinetics of nanoporous thiol–ene PEG hydrogels increased with increasing irradiation intensity, indicating faster percolation at higher light intensities based on the evolution of the storage modulus (G′). In each case, the light was turned on at $t$ = 45 s.

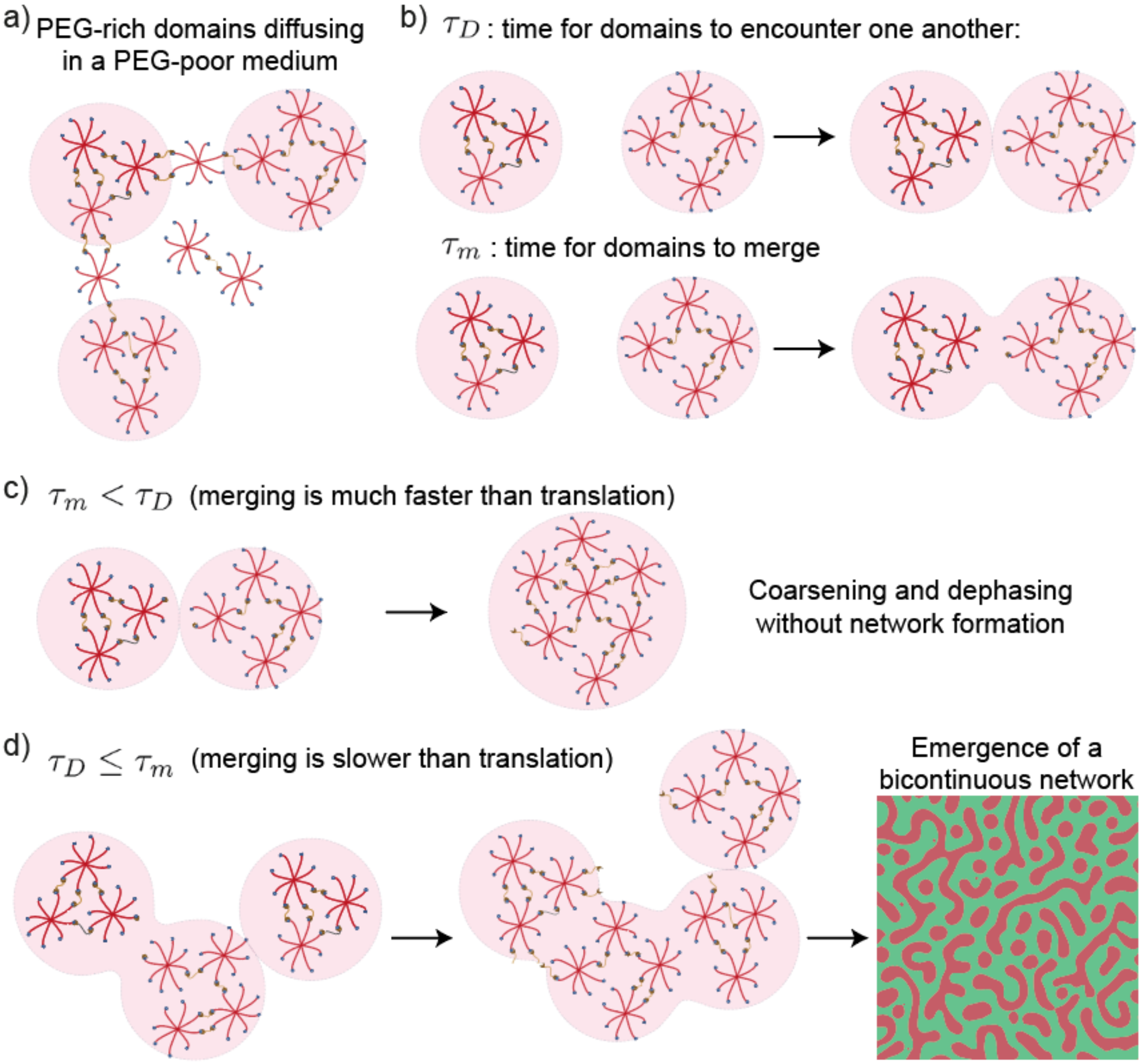


**Figure S3**. Schematic representation of the physical model of kinetic control of bicontinuous network formation. a) We assume that at the onset of phase separation PEG-rich domains (increased local concentration of PEG) form, which are suspended in a surrounding medium of PEG-poor solution. We assume cross-linking occurs within the domains and not in the surrounding solution. These domains are approximately spherical and can translocate via Brownian motion and can merge when they encounter another domain up to the point that the cross-linking density in the domains prevents further merging. b) There are two timescales that govern the evolution of this system during photopolymerization: $\tau_D$ describes the time for domains to encounter one another and $\tau_m$ describes the time for domains in contact to merge. c) If $\tau_m < \tau_D$, that is if merging is faster than translation, the domains can merge to form a larger, spherical domain. This process ultimately leads to coarsening and dephasing without the formation of a bicontinuous network. d) If $\tau_D \leq \tau_m$, that is if merging is slower than translation, additional domains will add to the merging domains forming elongated and anisotropic structures with characteristic length scale, ξ. This process ultimately leads to the emergence of a bicontinuous network. The balance between $\tau_m$ and $\tau_D$, is governed by system parameters including the rate of polymerization and solution viscosity, which provide handles to control the process.

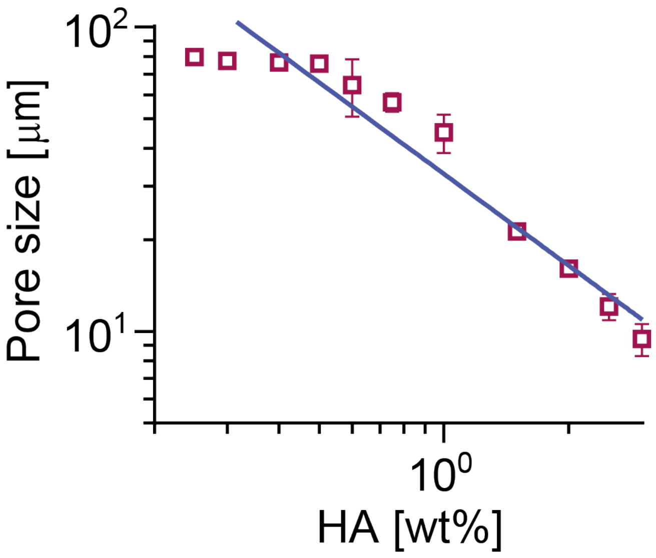


**Figure S4.** Pore size scaled with solution viscosity, assuming that it is proportional to HA concentration in the hydrogel precursor solution.
